# Spectral Interferences Impede the High-Resolution Mass Analysis of Recombinant Adeno-Associated Viruses

**DOI:** 10.1101/2022.08.27.505551

**Authors:** Victor Yin, Paul W.A. Devine, Janet C. Saunders, Alistair Hines, Sam Shepherd, Marcin Dembek, Claire L. Dobson, Joost Snijder, Nicholas J. Bond, Albert J.R. Heck

## Abstract

Recombinant adeno-associated viruses (rAAVs) are the leading platform for *in vivo* delivery of gene therapies, with several already approved for clinical use. However, the heterogeneity and structural complexity of these viral particles render them challenging targets to characterize. Orbitrap-based native mass spectrometry (MS) is a method capable of directly characterizing intact megadalton protein assemblies. Here we used such an approach to characterize four different preparations of rAAV8 (two empty and two filled) differing in both their transgene and relative capsid protein isoform (i.e. VP1, VP2 and VP3) content. Interestingly, in native MS measurements of these samples, we observe complicated, unusual, and dramatically different spectral appearances between the four rAAV preparations that cannot be rationalized or interpreted using conventional approaches (i.e. charge state deconvolution). By combining high-resolution native MS, single particle charge detection MS, and spectral simulations, we reveal that these unexpected features result from a combination of stochastic assembly-induced heterogeneity and divergent gas phase charging behaviour between the four rAAV preparations. Our results stress the often-neglected heterogeneity of rAAVs, but also highlight the pitfalls of standard high-resolution mass analysis for such particles. Finally, we show that charge detection MS and spectral simulations can be used to tackle these challenges.

## Introduction

Adeno-associated viruses (AAVs) are small, non-pathogenic, single-stranded DNA viruses that are incapable of replication in the absence of a helper virus.^1^ These favourable properties have resulted in increasing interest in the use of recombinant adeno-associated viruses (rAAVs) as a gene therapy vector due to the capability to pack rAAVs with a therapeutic gene of interest (GOI).^2–6^ A plethora of natural AAV serotypes are known, with a broad range of tropisms for different tissues types.^7^ Significant improvements have been made to these natural variants through capsid modifications, enabling targeting of specific cells types and reduced capsid-mediated immunogenicity.^8,9^

Although rather small (ca. 25 nm in diameter), the capsids of AAVs are heterogeneous and are constructed from three capsid proteins: VP1, VP2, and VP3. These three viral proteins (VPs) are produced from alternate initiation sites of the same gene, and thus share a large portion of the same amino acid sequence (**Figure 1A**).^10^ All three VP isoforms contain the full C-terminal VP3 sequence, which forms the primary structural component of the AAV capsid. VP1 and VP2 diverge from VP3 by the presence of extensions on the N-terminus of the protein, with the entirety of the VP2 sequence also contained in the VP1 sequence. Although the presence of VP1 and VP2 are not required to form the AAV capsid^11^, the VP1 and VP2 N-terminal extensions play critical roles in nuclear localization and genome release.^12^ Each icosahedral AAV capsid is composed of a total number of 60 copies of VP1, VP2, or VP3. Typical AAV preparations contain the three VPs in a ratio of approximately 5:5:50 (VP1:VP2:VP3), but can vary depending on serotype, expression system, and purification conditions.^13,14^

**Figure 1.**
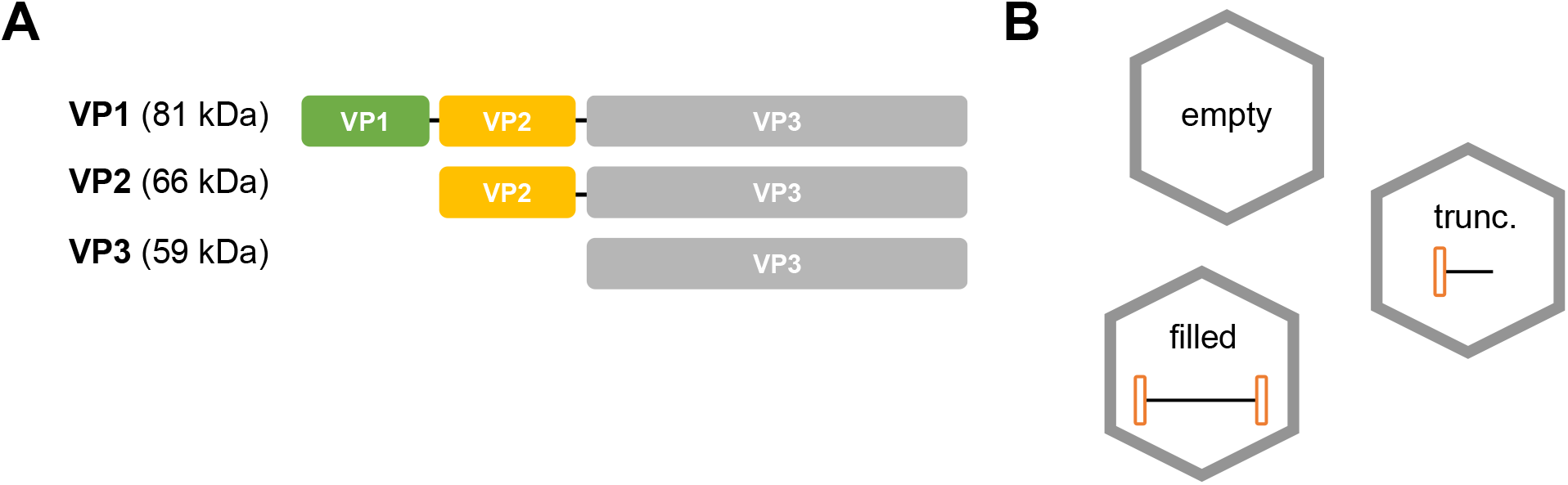
Compositional Heterogeneity of rAAVs. (A) AAV capsids are comprised of 60 capsid proteins of three different isoforms: VP1, VP2, and VP3. While the core capsid structure is formed by the shared VP3 region, the different N-terminal extensions of VP1 and VP2 add a degree of heterogeneity to each assembled capsid. (B) Transgene incorporation can vary within an rAAV preparation. Some particles may be empty, while others will contain the transgene. A fraction of particles may also contain incorrect (e.g. truncated or elongated) transgene, or even production host cell DNA.^**22**^

In parallel to the compositional heterogeneity present in the viral capsid itself, therapeutic rAAVs also contain a transgene component, further increasing the complexity and heterogeneity (**Figure 1B**). Current expression methods for rAAVs can yield up to 90% empty capsids (i.e. only 10% of particles actually contain DNA), depending highly on serotype, production and purification techniques.^15–17^ Attempts to package genomes larger than ca. 5 kb can also result in truncations of the packaged ssDNA, adding a further layer of heterogeneity.^18^ Other factors, such as formation of deleterious secondary structure within the DNA strand, can also negatively impact the fidelity of the packaged therapeutic GOI.^19,20^ Evidently, the integrity and packing efficiency of the transgene is critical for the function of rAAVs as a gene delivery vehicle.^21^

The complexity inherent to rAAVs results in a wide array of attributes that must be monitored, posing a substantial analytical challenge. Due to the breadth of these characteristics, multiple assays using a variety of analytical techniques are typically combined to obtain a comprehensive view of an rAAV preparation.^23^ For example, the ratio of the three VP isoforms in a preparation is commonly determined by disassembling the particles followed by electrophoretic analysis.^24–26^ While sufficient at resolving the 3 VP isoforms, this approach is invariant to the level of transgene incorporation, nor does it provide information on capsid integrity. On the other hand, the proportion of empty and genome-filled particles (“E/F content”) can be measured by separating the two populations using centrifugal^27^ or ion exchange chromatographic^28,29^ methods, or visualized by electron microscopy.^30,31^ While able to distinguish the presence or absence of DNA within an rAAV, these techniques typically do not determine any other transgene characteristics (e.g. identity, integrity), nor is any information obtained on the capsid VP compositions.

One property that is conveniently shared between these diverse rAAV attributes is that changes in any of these characteristics can be directly mapped to a change in mass of the rAAV. Therefore, determining the mass of rAAVs (and/or its constituent components) allows for determination of multiple rAAV attributes simultaneously. Orbitrap mass spectrometry (MS), with its high resolving power, is an analytical technique that is well-suited for this task.^32,33^ The high-resolution and high mass capabilities of Orbitrap mass analyzers allow for measurements of intact rAAVs and other large macromolecular assemblies in the megadalton regime.^34^ Moreover, different Orbitrap MS acquisition strategies allow for determination of different rAAV attributes, e.g. VP1/2/3 ratios^35^, capsid integrity^36^, or E/F content.^37^

Here, we present the combination of Orbitrap native MS-based measurements to deeply characterize several rAAV preparations, using the AAV8 serotype as a representative system. We demonstrate the excellent agreement between attributes determined *via* Orbitrap MS and conventional analytical methods, while simultaneously highlighting the deeper level of information obtained by Orbitrap native MS, such as stochastic heterogeneity of VP ratios between the capsids of a preparation, and integrity of the transgene. Importantly, we provide key practical considerations and pitfalls when performing and interpreting results from high-resolution native MS measurements of these heterogeneous complexes. Our results emphasize the utility of native MS approaches for characterizing rAAV attributes, thereby improving the toolbox available to the field for development of these highly complex gene therapy vehicles.

## Results

### rAAV Samples

For this study, we investigated four different preparations of rAAV8 (**Table 1**). These differ in both their transgene and relative VP isoform content, but were all produced from identical source plasmids. Samples S.E-0 and S.E-1 were produced in the absence of a transgene-containing plasmid (i.e. by double transfection), generating empty rAAV capsids. Samples S.F-0 and S.F-1 were produced *via* triple transfection, yielding rAAV preparations where a proportion of produced capsids contain a ZsGreen transgene.

**Table 1.** Attributes of the four rAAV8 preparations used in this work. The empty to full (E/F) content of each preparation was initially assessed by ion exchange chromatography (IEC), while the relative monomer ratios of each VP isoform were assayed by capillary gel electrophoresis (CGE).

| Sample Name | Serotype | Transfection Format | Transgene |  | VP Content |  |  |
| --- | --- | --- | --- | --- | --- | --- | --- |
|  |  |  | Identity | %Full | %VP1 | %VP2 | %VP3 |
| S.E-0 | AAV8 | Double | / | 0 | 8 | 12 | 80 |
| S.E-1 | AAV8 | Double | / | 0 | 7 | 12 | 82 |
| S.F-0 | AAV8 | Triple | ZsGreen | 20 | 8 | 23 | 68 |
| S.F-1 | AAV8 | Triple | ZsGreen | 16 | 10 | 25 | 65 |

### Characterizing rAAV Integrity and Transgene Packing Efficiency

To verify the integrity of each rAAV preparation, we first imaged each sample using negative stain transmission electron microscopy (nsTEM) (**Figure 2**). Visual inspection of the nsTEM images reveal that intact particles of ca. 25-30 nm are observed in all preparations, consistent with the expected rAAV diameter.^38^ No particles corresponding to partially assembled capsids were confidently identified in the images. As well as reporting on capsid integrity, nsTEM is also used to assess the E/F content of an rAAV preparation, as filled particles are often visually distinguishable from their empty counterparts.^31,39,40^ Oddly, the nsTEM results indicate the presence of a substantial proportion of filled capsids even in S.E-0 and S.E-1 (96% and 68%, respectively), despite being produced under conditions where no transgene incorporation is possible (**Figure 2A,B**). Moreover, the E/F content for S.F-0 and S.F-1 (50% and 37%, respectively) are in rather poor agreement between nsTEM and IEC, with nsTEM reporting elevated levels of genome incorporation (**Figure 2C,D)**.

**Figure 2.**
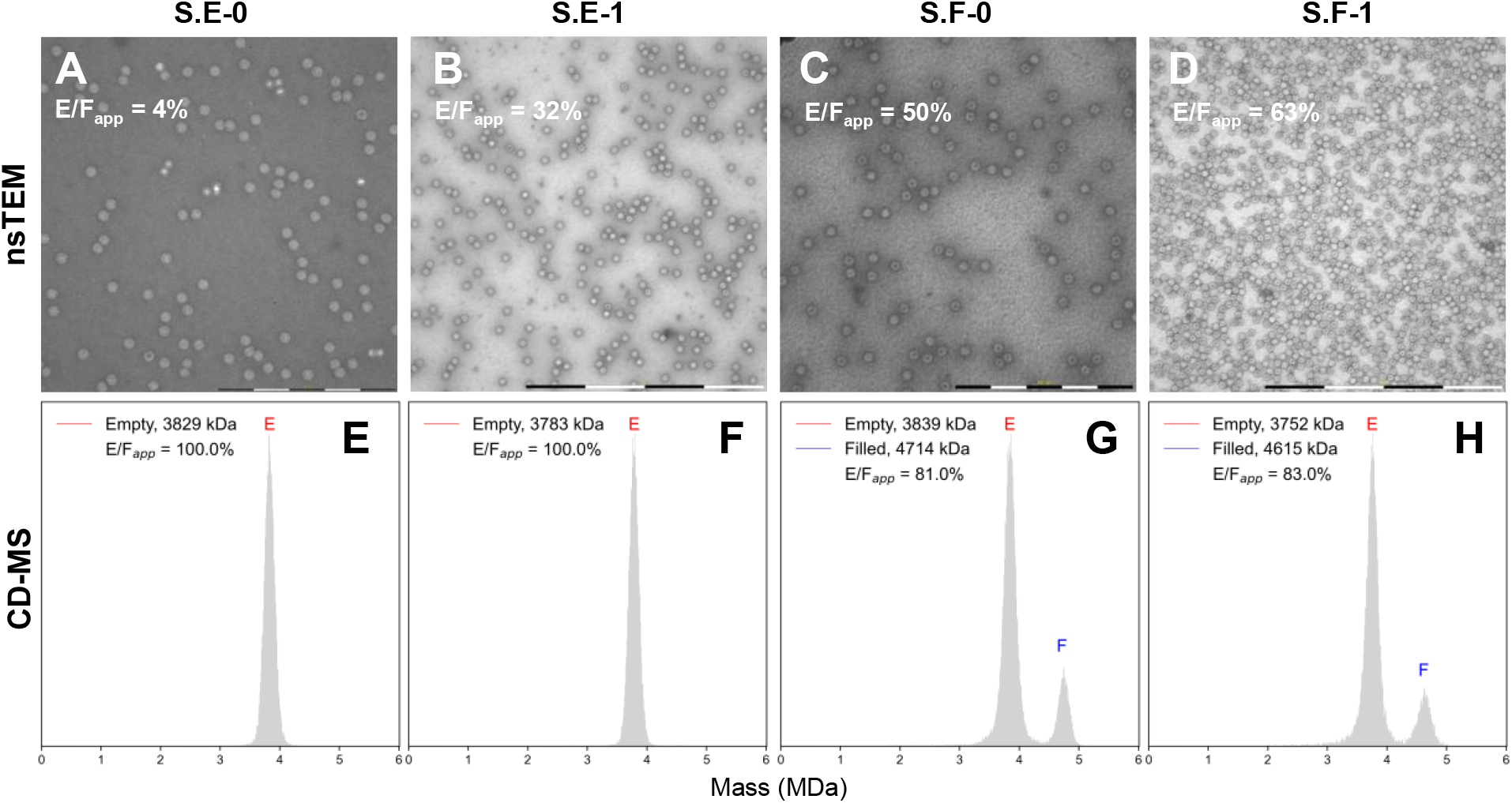
nsTEM and CD-MS of rAAV preparations. (A-D): Representative nsTEM images of each rAAV preparation. No evidence of partially assembled or aggregated particles were observed. Particles with internal staining, normally classified as filled rAAVs, are observed in all rAAV preparations. (E-H): CD-MS mass histograms of each rAAV preparation. Peak integration of the fully-resolved empty and filled capsid populations yield the E/F content. In contrast to the nsTEM data, the CD-MS histograms provide E/F content of each rAAV preparation in good agreement with those obtained by ion exchange chromatography (IEC). Scale bars for nsTEM: A+C, 500 nm; B+D, 1000 nm.

Inconsistencies between the E/F content derived from nsTEM and orthogonal analytical methods have been previously reported, and may arise from factors such as inhomogeneity in staining efficiency.^28,41^ This effect commonly leads to an overestimation of filled particles due to erroneous (filled) particle classification. To resolve these discrepancies, we employed Orbitrap-based single particle charge detection mass spectrometry (CD-MS) to orthogonally determine the E/F content of each rAAV sample (**Figure 2E-H**). CD-MS simultaneously measures both the mass-to-charge (*m/z*) and charge (*z*) of an individual ion in the gas phase, allowing direct mass determination of a given ion by the product of these two values.^36,42^ As Orbitrap UHMR mass spectrometers can efficiently measure masses in the MDa regime^34^, the integrity of the rAAV capsid, as well as its associated transgene, can be determined *via* CD-MS from their characteristic masses using minute amounts of sample.^43^ The relative ion abundances of empty and filled capsids detected by CD-MS have previously shown excellent quantitative accuracy across preparations.^37^

As expected from the double transfection method used to produce them, the CD-MS histograms of the S.E-0 and S.E-1 AAV8 samples (**Figure 2E-F**) both show a single mass distribution centered at ca. 3.8 MDa, corresponding to the expected mass of empty rAAV capsids. The CD-MS histograms of S.F-0 and S.F-1, by comparison, both have a second distribution at higher mass that is absent in S.E-0 and S.E-1, corresponding to the filled capsid population (**Figure 2G,H**). The E/F content obtained by CD-MS peak integration (ca. 19% and 17%, respectively), show excellent agreement with the results from IEC (20% and 16%, respectively). For additional confirmation, we also performed analytical band centrifugation (ABC) on S.F-1, with a comparable E/F ratio (17%) to both IEC and CD-MS obtained (**Figure S1**).^44^ These results support the ability of CD-MS to accurately measure E/F content, but also suggest that caution should be taken when interpreting E/F content obtained from nsTEM.^37^

Aside from low sample consumption, another advantage of CD-MS over other analytical methods of rAAV characterization is the ability to determine the E/F ratio and transgene integrity simultaneously within the same measurement, as any truncations or extensions in the transgene will result in differences in the mass of the particles. In the cases of both S.F-0 and S.F-1, the mass difference of the filled and empty capsid populations (ca. 870 kDa) agrees reasonably with the sequence-predicted molecular weight of the ZsGreen transgene itself (ca. 820 kDa, 2.6 kbp). The minor discrepancy in mass may arise from incomplete ion desolvation or small amounts of counter-ions packaged alongside the transgene. Additionally, the filled capsid populations exhibit peak widths comparable to that of the empty capsids, demonstrating that the transgene packaging is rather monodisperse. Overall, the CD-MS results support the integrity of transgene packaging in these rAAV samples, with no evidence of truncated gene products (which would manifest as intermediate masses between the empty and filled populations) in these preparations.^37^ These results highlight the utility of CD-MS measurements to probe transgene-related rAAV attributes.

### Interrogating VP Distributions Within Intact rAAV Capsids by High Resolution Native MS

Contemporary structural biology methods (e.g. single particle cryoTEM) are unable to distinguish the different VP isoforms present in a capsid due to the low occupancy and conformational heterogeneity of the VP1 and VP2 N-terminal components. Thus, in the absence of experimental methods capable of directly probing isoform distributions within an assembled capsid, it has long been assumed that intact AAV capsid particles must be monodisperse, such that every individual rAAV particle contains an identical copy number of each VP species.^45^ The relative ratios of VP1, VP2, and VP3 in an rAAV preparation can thus be validly determined by disassembling the capsids and subsequently quantifying the abundance of each VP isoform by e.g. electrophoresis or LC-MS.^24–26^

High-resolution native MS was recently shown to be capable of directly monitoring the distribution of VP isoforms within intact capsids.^35,46^ In a high-resolution native MS experiment, standard charge state inference strategies are utilized to indirectly assign charge state values to each species.^47,48^ The primary advantage of this approach is that a substantially higher mass resolution to CD-MS can be obtained as there is essentially no uncertainty in charge determination - with the caveat that the mass spectra must be sufficiently well-resolved for charge states to be accurately inferred. In cases where this requirement cannot be fulfilled, CD-MS out-performs standard native MS in mass determination ability.^49^ To simplify the terminology used herein, the more conventional native MS experiments will be refered to simply as “native MS”, although the single particle CD-MS measurements described above can be formally classified as a sub-type of native MS experiment.^33^

### Spectral Interferences Lead to Unusual Native Mass Spectral Appearances for rAAVs

High-resolution native mass spectra of each rAAV sample were acquired and are depicted in **Figure 3**. For simplicity, the subsequent discussion will be focused on the majority capsid population (i.e. empty) of each of the four studied rAAV preparations. Compared to the mass spectrum of a theoretical, monodisperse rAAV particle (e.g. VP3 only, **Figure 3A**), each experimental rAAV spectrum exhibits substantially more peaks in the *m/z* domain (**Figure 3B-D**), reflecting the heterogeneity of the underlying capsid VP compositions. Similar spectral appearances have been reported previously for other rAAVs.^35,46^ Early reports attributed this peak multiplicity primarily to the presence of between three and six different capsid VP stoichiometries, as there are approximately 3 times more peaks than expected in the *m/z* domain.^46^ Of the samples analyzed here, S.E-0 and S.F-0 appear to be consistent with this interpretation, as approximately three times the amount of peaks are observed in a seemingly well-resolved charge state distribution (**Figure 3B,D**). However, the native mass spectra of S.E-1 and S.F-1 display different spectral appearances over the same measured *m/z* window (**Figure 3C,E**). While a similar high-multiplicity distribution to S.E-0 and S.F-0 is also present in S.E-1 at ca. 22,500 *m/z*, a second “low-resolution” distribution at ca. 24,000 *m/z* is observed in the same spectrum (**Figure 3C**). A similar distribution is also present in S.F-1, where it appears to comprise most, if not all, of the mass spectrum (**Figure 3E**). The unusual appearance of S.F-1 is especially surprising given that S.F-0 and S.F-1 contain very similar levels of each VP isoform, yet seem to exhibit dramatically different spectral appearances (**Table 1**).

**Figure 3.**
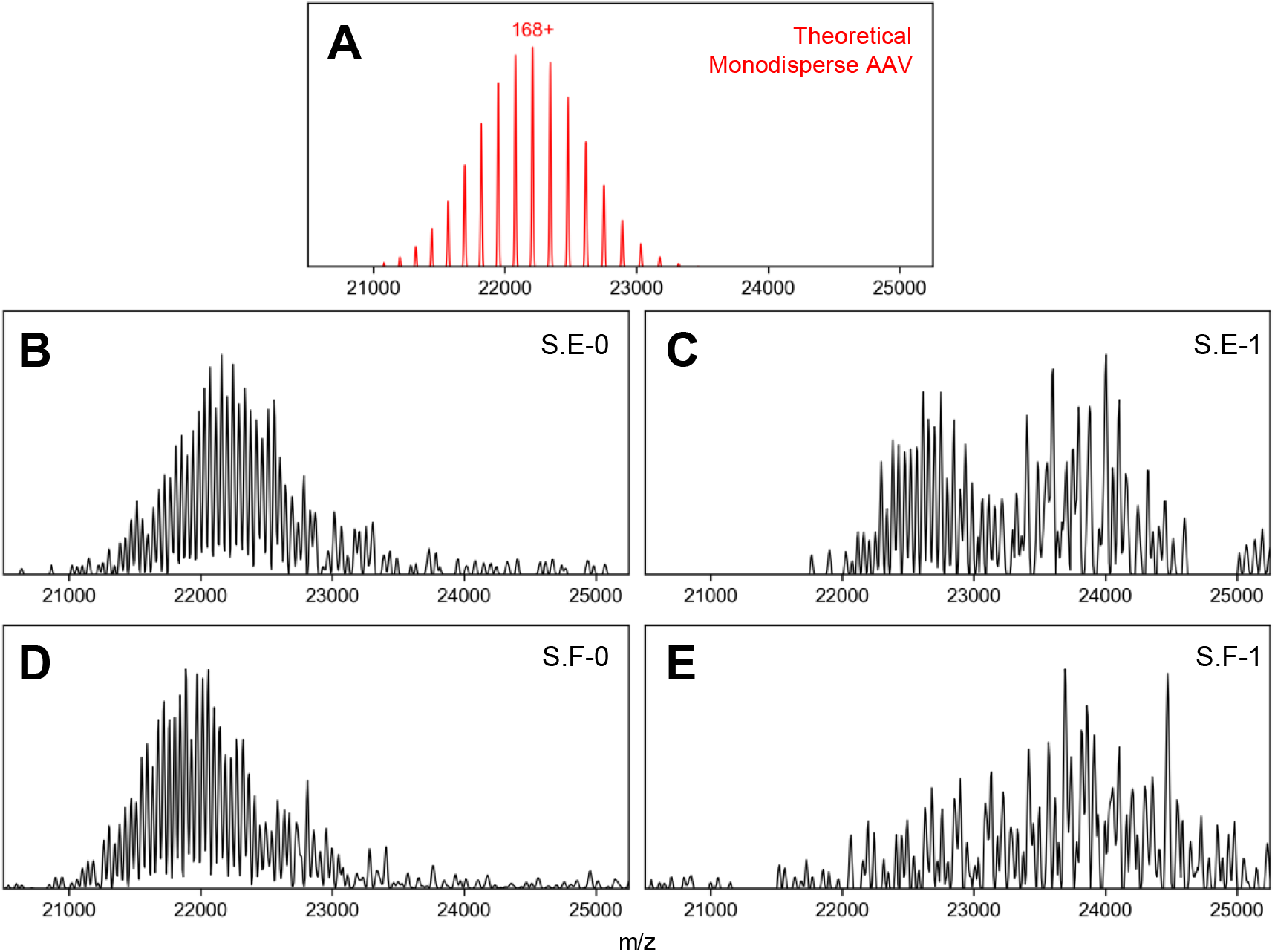
High-resolution native MS of four different rAAV8s. (A) Simulated native mass spectrum of a monodisperse AAV particle (e.g. (VP3)_60_). Only a highly resolved single charge state distribution is observed. (B-E): Experimental high-resolution native mass spectra of each rAAV8 preparation. For each spectrum only the ion signals corresponding to the empty rAAVs are shown and in each spectrum the same *m/z* window is depicted.

### Spectral Simulations of Stochastic Assembly

Rather than dismissing the unusual spectral appearances of S.E-1 and S.F-1 as a consequence of e.g. poor sample quality or experimental artefacts, we instead hypothesized that their unusual spectral features arise from an intrinsic property of the rAAV capsids themselves – specifically in the heterogeneity of their VP compositions. Recently, Wörner *et al*. proposed that rAAV particles assemble with an entirely statistical probability distribution of the VP isoforms (i.e. stochastically).^35^ In this model, instead of a monodisperse population (or near-monodisperse, such as the ~ 3 stoichiometries seemingly observed above in **Figure 3**), a given rAAV preparation can be described as a mixture of all the 1,891 combinatorially possible capsid compositions, with the relative abundance of each composition dependent solely on the relative abundances of each VP isoform in the particle-producing host cell.

Counterintuitively, the work of Wörner and coworkers predicts that native mass spectra of rAAVs will not directly reflect all 1,891 stoichiometries, which would be near-impossible to resolve due to the immense expected spectral complexity. Instead, due to the non-zero peak widths of mass spectral signals, peaks that coincidentally have similar *m/z* positions (but originate from different charge states of different capsid compositions) are expected to coalesce into a single apparent peak. When considered over all *m/z* values, this phenomenon effectively produces “spectral interference patterns” of signals in the native mass spectra, whose appearance is dependent on the underlying capsid compositions, but displays substantially fewer resolved peaks than expected *a priori*. Critically, the appearance of these interference patterns is expected to be exquisitely sensitive to both the masses (i.e. *m/z* position) and relative abundances of each VP isoform.

To determine whether these effects are the root cause of the abnormal spectral features displayed in **Figure 3**, we performed spectral simulations of each of the native mass spectra by applying the stochastic assembly model proposed by Wörner *et al*., using the CGE-derived VP isoform ratios as input (**Figure 4**).^35^ In brief, the experimental native mass spectra are assumed to be a linear combination of individual native mass spectra from all 1,891 possible capsid compositions, weighted by their statistically-predicted abundances. Peaks from different compositions, but exhibiting similar *m/z* values, are summed into a single peak to account for the experimentally-observed non-zero peak widths. As depicted in **Figure 4**, the native mass spectra simulated in this manner agree remarkably well with the experimental spectra, offering insight into the unusual spectral features, and confirming that experimental rAAV spectra can be validly represented by a stochastic assembly model.

**Figure 4.**
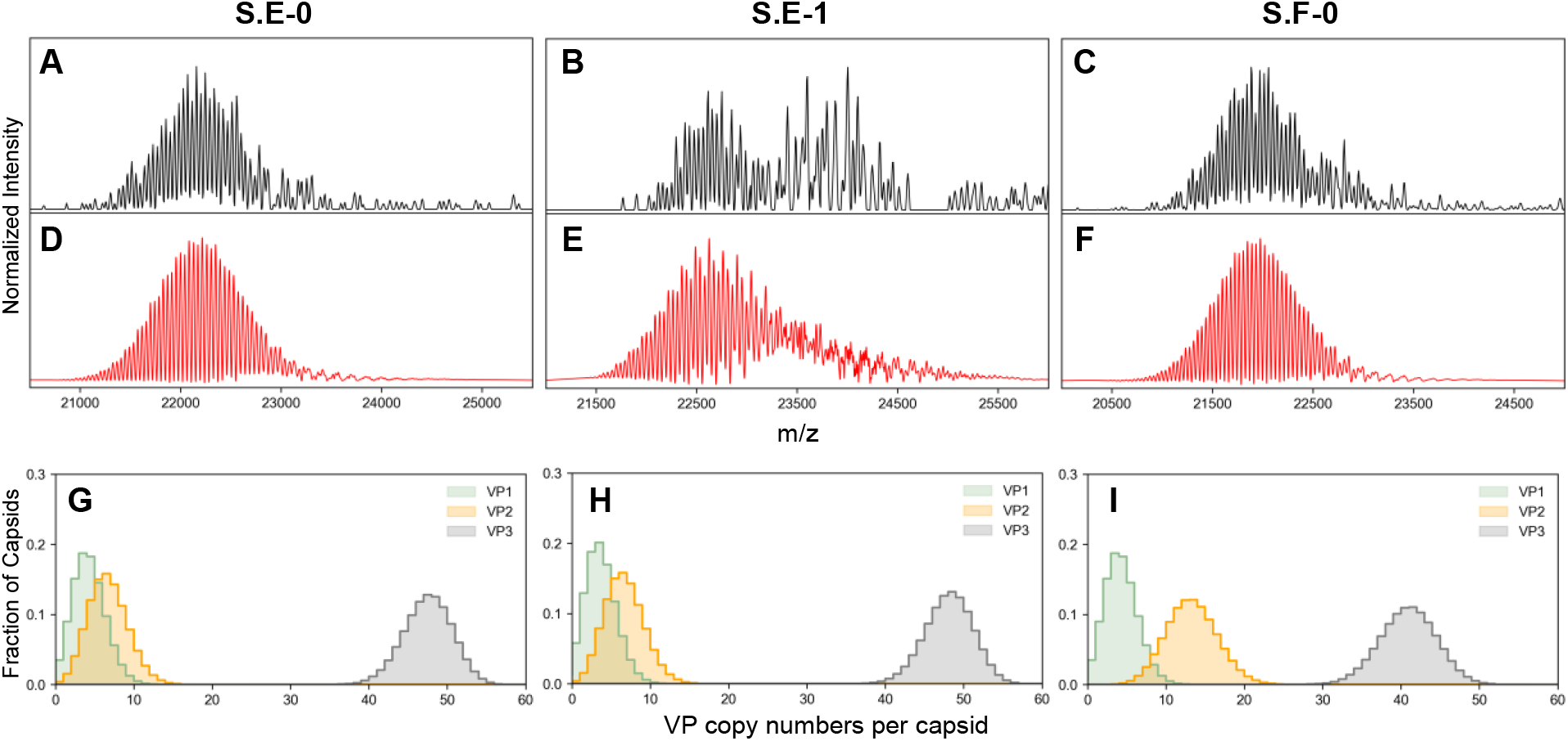
Comparison of experimental data and spectral simulations of S.E-0, S.E-1, and S.F-0. (A-C): Experimental native mass spectra of each rAAV preparation, as shown in Figure 3. (D-F): Simulated native mass spectra of each rAAV preparation following the stochastic assembly model proposed by Wörner *et al*.^**35**^ Very good agreement between the experimental and simulated mass spectra is observed. (G-I): Probability histograms of the VP isoforms in each rAAV preparation extracted from the spectral simulations, highlighting the heterogeneity within each capsid population.

For S.E-0 and S.F-0, the simulated spectra correctly recapitulate both the high multiplicity and peak positions seen in the experimental spectra (**Figure 4D,F**). These spectral appearances are comparable to those reported by Wörner *et al*., where similar results were observed across a variety of AAV serotypes obtained from independent sources.^35^ In contrast, a different spectral appearance is observed here in the simulated spectra of S.E-1 (**Figure 4E**). Although a seemingly highly resolved region is observed in the simulated spectrum between ca. 22,000 and 23,500 *m/z*, this collapses into a poorly-resolved, heterogeneous region from ca. 23,500 to 24,500 *m/z*. Interestingly, this poorly-resolved region in the simulated spectrum coincides extremely well with the “low resolution” region seen in the experimental spectrum of S.E-1 (**Figure 4B)**. In other words, these spectral simulations reveal that the second, poorly-resolved distribution is not an experimental artefact, but is an intrinsic, expected feature of the experimental rAAV native mass spectrum.

The unusual loss of apparent resolution in this *m/z* range can be rationalized by the existence of separate interference windows in different regions of the mass spectrum. As described above, peaks in the native mass spectra of rAAVs can be attributed to a constructive interference pattern of constituent native mass spectra corresponding to each unique capsid distribution. The predominant contributions that give rise to the “high multiplicity” region at ca. 21,000 *m/z* are that the substitutions of either 3 VP3 → VP2 or 1 VP3 → VP1 (ca. 21 kDa) produce a shift in the *m/z* domain that is extremely similar to the spacing between adjacent charge states of a single capsid distribution at these charge values.^35^ For example, assuming an rAAV mass of 3.8 MDa and a charge of 182+ (ca. 21,000 *m/z*):

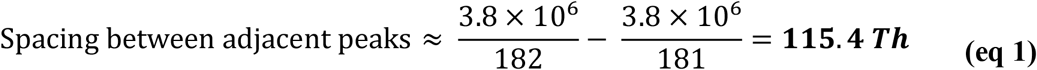

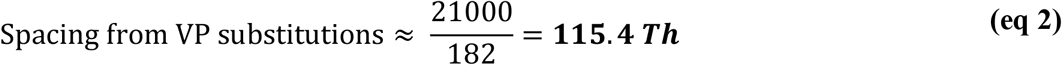

By comparison, at higher *m/z* values (= lower charge; ~ 157+), these two shifts dephase and constructive interference becomes weaker, e.g.:

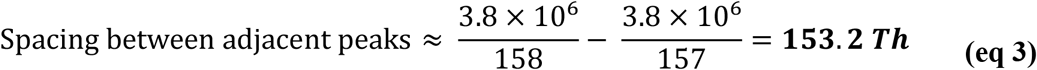

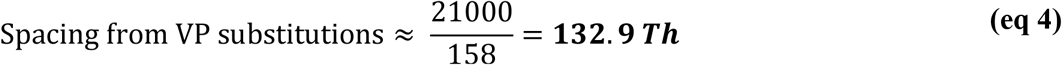

In S.E-0 and S.F-0, it appears that the majority of the rAAV spectral signal is concentrated in the *m/z* window where constructive interference is dominant (**Figure 4D,F**). Conversely, in S.E-1 the signals are split between two windows: one where constructive interference is dominant, and another where the signals are dephased (**Figure 4E**). The native mass spectra of S.F-1, whose appearance is particularly unusual (**Figure 5A**), can thus be attributed to a particularly extreme case of this phenomenon, where essentially all of the spectral signal reside within an *m/z* window hampered by strong interference (**Figure 5B**).

**Figure 5.**
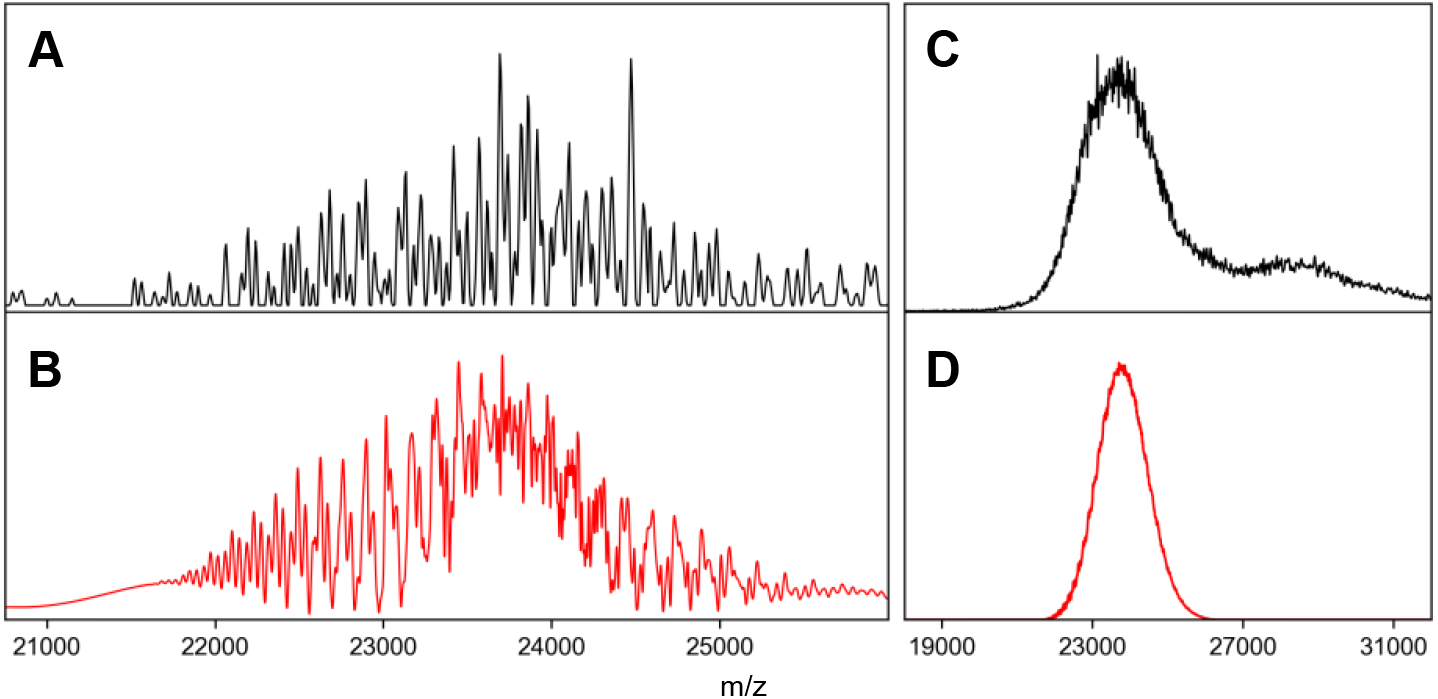
Spectral simulations rationalize the abnormal spectral features observed for S.F-1 AAV. (A) Experimental native mass spectrum of S.F-1 collected using time-domain averaging. (B) Simulated native mass spectrum of S.F-1 using the stochastic assembly model mimicking time-domain averaging (i.e. baseline correction), recapitulating quite well the complex signal ensemble observed in (A). (C) Experimental native mass spectrum of S.F-1 collected using frequency-domain averaging confirm that signal corresponding to intact rAAVs can be obtained. (D) Simulated native mass spectrum of S.F-1 using the stochastic assembly model, mimicking frequency-domain averaging.

It is important to clearly emphasize that due to these stochastic assembly effects, AAV spectra such as those of S.F-1 will not present clearly defined apparent charge state distributions, and can instead produce highly complex / amorphous spectral features. These effects are independent of e.g. low S/N. To demonstrate this, native MS data of S.F-1 was also recorded with transient averaging disabled, and instead averaged in the frequency domain (i.e. post-FT averaging, **Figure 5C**). Under these instrument settings, an intense, albeit unresolved, signal corresponding to the empty rAAV can be clearly observed, with a similar smaller signal at ca. 28,000 *m/z* corresponding to the filled rAAV population. An identical broad signal can be reproduced *via* spectral simulations, if modified to mimic the frequency-domain averaging performed experimentally (**Figure 5D**). These observations show that care must be exercised when collecting or interpreting transient-averaged AAV spectral data, as their spectral appearances and apparent intensities will be extremely sensitive to the stochastic assembly effects described here. These effects greatly complicate the extraction of accurate mass information from high-resolution AAV native mass spectra, even when a portion of the spectra appear “charge-state resolved” (such as the data presented in **Figure 4A-C**). However, by using the spectrum simulation strategy described here, mass information of all particles can regardless be obtained, alongside e.g. abundance of all particles with given VP isoform ratios (**Figure 4G-I**).

### Charging Behaviour Dramatically Affects the Observed Spectral Features in Different rAAV Preparations

We were at first quite surprised that such similar rAAV preparations exhibit such drastically different mass spectral appearances. While the average mass of each sample is expected to vary slightly due to the different relative ratios of VP isoforms, the contribution of this effect alone is expected to be fairly minor. More impactfully, the positions of the rAAV spectral signals in the mass spectrum is strongly dependent on their obtained charge following electrospray ionization (ESI). The charge states for each rAAV is challenging to accurately ascertain from the native mass spectra (e.g. **Figure 3**) due to the stochastic assembly effects described above. To solve this issue, we revisited the CD-MS data presented in **Figure 2**. While in **Figure 2** only the final determined mass of each ion is depicted, the corresponding charge of each ion is directly measured as an intermediate step of the CD-MS method. Interrogating these measured charges in the two-dimensional CD-MS spectra (i.e. *z* vs. *m/z*) allow a direct assessment of the charge state distributions of each rAAV preparation (**Figure 6**).

**Figure 6.**
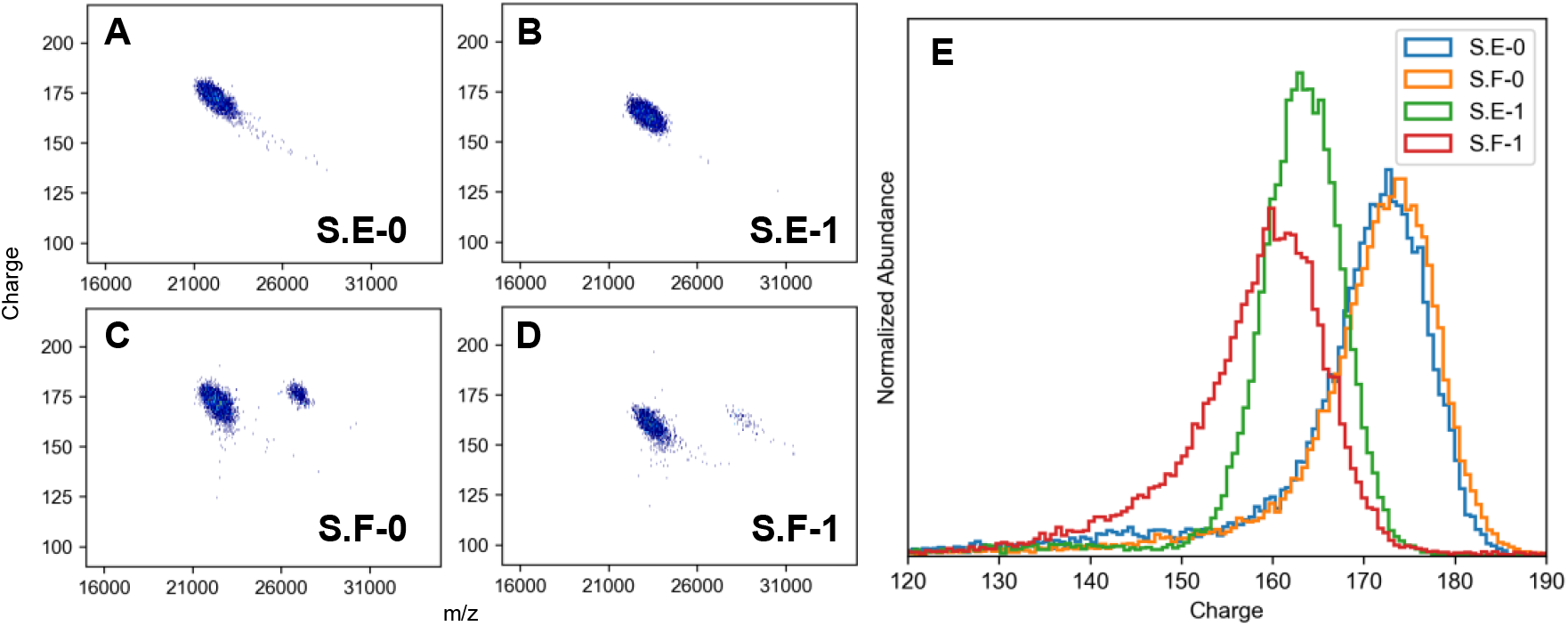
ESI charging behaviour of each rAAV preparation as determined by CD-MS. (A-D): 2D CD-MS histograms of each rAAV preparation. For S.F-0 and S.F-1, a second distribution at higher *m/z* is also observed, corresponding to the genome-filled rAAV. In both cases, the filled rAAVs possess very similar charge to their empty counterparts, despite being ca. 20% higher in mass.^**37,50**^ (E): Overlaid 1D charge histograms of each rAAV preparation.

The charging behaviour of each of the rAAV preparations revealed by CD-MS offers an explanation to their distinct native mass spectral appearances. S.E-0 and S.F-0 exhibit near-identical ESI charging, with an average charge of ca. 175^+^ (**Figure 6A,C,E**), and also show very similar appearances in the native mass spectra (**Figure 3B,D**). In contrast, S.F-1 carries a comparatively lower charge, centered at ca. 160^+^, and exhibits a high degree of dephasing in its native mass spectra (**Figure 3E**). S.E-1 carries an intermediate level of charge between two extremes, thus rationalizing the combination of both phased and dephased peaks seen in the native mass spectra (**Figure 3C**).

While the CD-MS data presented above pinpoints differential ESI charging effects as the primary source of the spectral appearances in **Figure 3**, the underlying cause of this behaviour remains somewhat nebulous. For approximately spherical protein complexes, the ESI charge states are predicted to closely follow the so-called Rayleigh limit of an equivalently sized water nanodroplet.^51,52^ For AAVs, this would correspond to an expected charge of 176^+^ (assuming a diameter of 25 nm^40^). While this model nicely correlates with the observed charges of S.E-0 and S.F-0 (**Figure 6A,C**), it overestimates the charges for S.E-1 and S.F-1, which are comparatively lower (**Figure 6B,D**). Unusually low ESI charging behaviour of other viral particles have been observed before. For example, a recent report by Williams and co-workers observed under-charging on the order of 30% for MS2 viral capsids, which was postulated to arise from conformational collapse of the nascent capsid ions within the electrospray droplet.^53^ A similar collapse thus may also be possible for the rAAV capsids studied here.

Further deviations from idealized (i.e. spherical) charging behaviour are likely expected in rAAVs due to the presence of the large VP1/2 N-terminal extensions (**Figure 1A**), which are in different abundances in each rAAV preparation (**Table 1**). Given that structural aspects of the VP1/2 extensions in AAVs are still poorly understood^45,54^, determining the precise relationship between rAAV structure and ESI behaviour is presently unfeasible. Nevertheless, our results clearly highlight the capacity for CD-MS to directly report on these changes in ionization behaviour.

## Discussion

The accurate characterization of rAAV attributes poses a substantial analytical challenge, but is a critical aspect to the application of rAAVs as possible therapeutics. Here, we report on the utility of Orbitrap native MS experiments to directly and simultaneously determine these characteristics that are challenging or unobtainable using other methods: single particle CD-MS allows for facile determination of E/F content and transgene integrity, while conventional native MS reports on the heterogeneous assemblies of the VP isoforms directly from the intact rAAV capsid. The work described here highlights the capacity of native MS experiments to provide detailed characteristics of rAAVs, and strongly supports its inclusion into the methodological toolbox of rAAV analysis techniques. Compared to other analytical methods, native MS has the advantage of low sample consumption (<10^11^ capsids per native MS measurement, ~ 10^9^ capsids for CD-MS), straightforward sample preparation, high sensitivity, and the capability to extract multiple key quality attributes simultaneously.

Our results demonstrate that the native spectral appearance of an rAAV is highly sensitive to its ESI charging behaviour, which can vary between preparations. It is important to emphasize that our observations here (that apparent spectral quality can decrease at lower charge values) runs contrary to the conventional wisdom in the field of native mass spectrometry, where lower charge values are expected to universally improve spectral quality *via* increasing of the spacing between adjacent peaks.^55,56^ Interestingly, this behaviour would imply that certain rAAV preparations, such as S.F-1 studied here, may not be expected to ever generate the “resolved” native mass spectra observed in S.E-0 and S.F-0 due to their intrinsic charging behaviour placing them in a poorly-interfering *m/z* window. While such results could be naively attributed to poor-quality data sets, our simulations confirm that these unusual spectral appearances are in fact intrinsic to the rAAV preparation, and can be fully rationalized. The spectral phenomena outlined in this work may at least partially explain the apparent inability of existing reports to obtain “high-resolution” native mass spectra of rAAVs.^57^ In this context, Orbitrap-based CD-MS offers a convenient method to determine the charging behaviour of an rAAV preparation, and would allow predicting *a priori* the native MS spectral appearances.

It is interesting to note that a statistical assembly model of capsid formation is mandatory to adequately explain all the appearances of the rAAV native mass spectra we observe (e.g. **Figure 3**). This stands somewhat in contrast to previous reports of high-resolution rAAV native mass spectra, which could be reasonably (albeit incorrectly) interpreted as a simple mixture of only a small number of capsid stoichiometries due to their classically well-resolved appearances (i.e. analogous to S.E-0 and S.F-0 here).^35,46^ Thus, our native MS data here provides the strongest evidence to date that rAAV capsid assembly must be a divergent, fully stochastic process. We stress that this stochastic heterogeneity is a generally over-looked attribute of rAAV capsids since it is essentially undetectable using other, contemporary methods of rAAV characterization. High-resolution native MS stands unique in its capability to detect this phenonmenon.^35^ The potential impact of this natural heterogeneity on the efficacy of rAAVs remains under-studied. For example, in a hypothetical rAAV preparation with a 0.05/0.05/0.9 ratio of VP1/2/3, we calculate that more than 9% of these capsids will be completely lacking in either VP1 or VP2 (i.e. likely functionally impaired). The native MS methodology outlined here sets the stage for future work in characterizing this important class of gene delivery vectors.

## Supporting Information

ABC analysis of AAV preparations, tabulation of key experimental MS parameters.

## Acknowledgements

We thank the members of the Heck laboratory for general support, especially Arjan Barendregt for technical expertise. This research received funding by the Netherlands Organization for Scientific Research (NWO) through the Spinoza Award SPI.2017.028 to AJRH. This project received funding from AstraZeneca.

## Author Contributions

Project conception: VY, NJB, CD, JS, and AJRH.

Sample production: MD, JCS nsTEM: PWAD

Native MS: VY

Supervision: NJB, CD, JS, AJRH

Writing: VY and AJRH with input from all authors

## Methods

### rAAV Expression and Purification

HEK Viral Production cells (ThermoFisher) were maintained in suspension in Freestyle F17 (ThermoFisher) chemically defined, serum-free media supplemented with 1xGlutamax (Gibco). The cells were cultured in 2L roller bottles (Greiner) at 37°C, 7% CO_2_ and 135 rpm agitation in a humidified atmosphere. The plasmids used were pAAV_zsGreen, pAAV_RC8, pAAV_Helper (produced in-house). Cells were seeded into 9 L medium at 0.5 × 10^6^ cells/mL in a wave bioreactor (Cytiva) on a Wave25 platform. 24 hours post-inoculation the culture was triple transfected using PEIMax (Polyplus) and three plasmids at a ratio of 2:1.5:1. To produce empty AAV particles, the culture were transfected identically except with the absence of the transgene plasmid (pAAV_zsGreen).

Each culture was harvested by batch centrifugation (3,800 x g for 15 min at 4°C) 72 hours post transfection to separate the cell fraction from the supernatant. The resulting cell pellet was freeze-thawed thrice and re-suspended to 10% (w/v) in a lysis buffer containing 5% (v/v) Polysorbate 80 and 20 U/mL benzonase (Merck). After 1h incubation at 37°C, the lysate was clarified by depth filtration followed by sequential filtration through 0.8 and 0.45µm filters. The clarified lysate was pooled with filtered supernatant and concentrated by TFF on an Ultracel Pellicon 2, 300 kDa MWCO cassette (Merck) before loading onto a pre-equlibrated POROS CaptureSelect AAVX affinity column (Thermo Fisher) at 1E+13 vg per mL of resin. After re-equlibration, AAV was eluted with a low-pH buffer and immediately neutralised with 1M Tris base. Neutralised eluate was concentrated and buffer exchanged into PBS using an Amicon Ultra, 100 kDa MWCO concentrator (Merck) before storage at −80°C.

### CGE

AAV samples were diluted to a concentration of 2E+12 vg/mL, using UltraPure water. Samples are then diluted with the Sciex IgG Purity/Heterogeneity kit sample buffer (Sciex, MA, USA) and beta-mercaptoethanol (BME) (Sigma), in a ratio of 10:9:1 (Sample:buffer:BME) resulting in a final concentration of 1E+12 vg/mL with 5% BME. Samples are heated for 10 minutes at 75 °C before being centrifuged for 30 seconds at 13,200 rpm, to cool and collect any condensation. The resulting AAV sample is then buffer exchanged three times into 1.7 mM SDS, 5% BME buffer exchange solution (Sciex, MA, USA), using 10 kDA MWCO filters (Vivaproducts Inc., MA, USA). 10 µL of the concentrated, buffer exchanged material is diluted to 100 µL, using UltraPure water, prior to loading onto a Sciex PA800 Plus instrument (Sciex, MA, USA). Instrument run conditions are as follows: sample injection, 5.0 kV for 60 seconds; separation, 15.0 kV at 25°C for 30 minutes; detection, 214 nm with a 10 nm bandwidth. Capillary used for analysis is a bare fused silica capillary with a 50 µm internal diameter, 30.2 cm total length (20.2 cm effective length), in a cartridge with 100 × 200 µm aperture. All data is processed using Empower (Empower, NY, USA).

### Analytical Band Centrifugation

Analytical band centrifugation experiments were performed following an adapted protocol.^44^ Briefly, analytical ultracentrifugation cells were fitted with band forming centrepieces with two sectors (Spin Analytical, USA) and 10 µL of AAV loaded into the sample reservoir. Samples were analyzed using a Beckman Optima system (Beckman Coulter, USA), with the following method parameters: equilibration period of 30 minutes at 0 rpm, at 20 °C. The material was then sedimented using a rotor speed of 20,000 rpm. There was a total of 150 scans measuring at 230 nm, with a scan frequency of 60 seconds at a resolution of 10 µm. The minimum and maximum radii for the experiment were 5.8 and 7.2, respectively. All data was analysed using an in-house Python script.

### Negative Stain Transmission Electron Microscopy

Carbon-coated copper 400 mesh grids (Ted Pella, CA, USA) were glow discharged for 1 minute using a PELCO easiGlow system (Ted Pella, CA, USA), prior to sample preparation. A 10 µL droplet of sample was pipetted onto parafilm and the prepared grids placed on top of the droplet, allowing adhering of the sample for 60 seconds. Excess sample was removed using Whatman filter paper, before washing the grid on two consecutive 20 µL droplets of ddH_2_O, before excess water was again removed by blotting against filter paper. Finally, the sample embedded grids were then stained by placing the grids on to a 20 µL droplet of 1% uranyl acetate for 30 seconds, where excess stain was then removed by blotting with filter paper. The prepared grids were then visualised and examined using a Vironova miniTEM system (Vironova, Sweden). Images were collected at a range of magnifications, with images for quantitation being captured at 1 and 1.5 µm field of view (FOV).

### Native MS

rAAV samples for native MS measurements were buffer exchanged into 75 mM ammonium acetate pH 7.5 by serial dilution using 50 kDa Amicon-0.5 MWCO centrifuge filter units (Sigma-Aldrich). For each sample, approximately 30 uL was centrifuged for 6 cycles of 10 minutes each at 9,000 G.

All native MS measurements were performed on an Orbitrap Q Exactive UHMR mass spectrometer (Thermo Fisher Scientific).^34^ For each measurement, approximately 1.5 uL of sample was loaded into a gold-coated borosilicate capillary (prepared in-house), and electrosprayed in positive ion mode. Xenon was used as collision gas in all measurements. A tabulation of typical instrument parameters for both conventional native MS and CD-MS can be found in **Table S1**. Unless otherwise noted, a transient time of 32 ms (with time-domain averaging enabled) was used for all conventional native MS measurements, while a transient time of 1024 ms (with time-domain averaging disabled) was used for all CD-MS measurements.

Samples for conventional native MS were typically analyzed without further dilution following buffer exchange. For CD-MS measurements, rAAV samples were diluted by ca. 100-fold prior to measurement. Ion injection times were manually attenuated to reach the single particle detection regime.

### MS Data Processing

Single particle CD-MS data was processed in Python as previously described using an intensity-to-charge calibration factor of 12.521.^36^ Simulations of the rAAV native mass spectra were performed following the model of Wörner et. al.^35^ In brief, the theoretical assembled rAAV capsid probability distribution is calculated *via* a stochastic assembly model. For any given VP1/2/3 monomer ratio, a simulated native mass spectrum for an rAAV preparation can thus be constructed by the probability-weighted average mass spectrum composed of the spectra from all 1,891 possible particle VP stoichiometries. Native mass spectra for each rAAV preparation were simulated in this manner using the CGE-derived VP1/2/3 monomer ratios as input. The charge states of the simulated mass spectra is a tunable parameter, and values were chosen to reflect the experimental data.

