## Supplemental Tables and Figures for "Spectral Interferences Impede the High-Resolution Mass Analysis of Recombinant Adeno-Associated Viruses"

This file contains:

**Supporting Figures**

Supporting Figure S1: Analytical band centrifugation of AAV preparation

**Supporting Tables**

Supporting Table S1: Key experimental parameters for native MS analysis

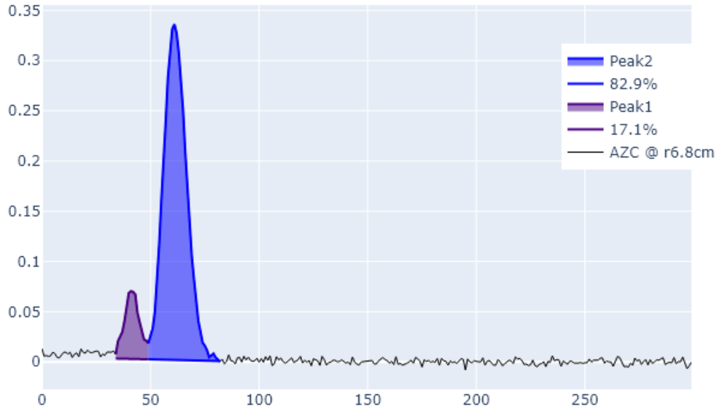


**Figure S1. Separation and Quantification of Empty and Filled S.F-1 by Analytical Band Centrifugation.** Two peaks are clearly separated of which the first one represent the filled AAV8 particles (17%) and the latter peak represents the empty particles (83%).

| | **Standard Native MS** | **CD-MS** |
|---|---|---|
| Capillary voltage (kV) | 1.30 | |
| *m/z* range | 10000 – 40000 | |
| Ion injection time (ms) | 30 | 10 - 1000 |
| Transient time (ms) | 32 | 1024 |
| Microscans | 10 | 1 |
| Averaging | 1000 | 0 |
| Noise threshold | 3.64 | 0 |
| In-source trapping voltage (V) | -75 | |
| HCD voltage (V) | 150 | |
| Trap gas setting | 4.0 | 1.0 – 2.5 |
| UHV readout (1e-10 mbar) | ca. 6 - 8 | ca. 1.4 to 4 |
| Collision gas | Xenon | |
| Injection flatapole (V) | 10 | |
| Inter-flatapole lens (V) | 10 | |
| Bent flatapole (V) | 4 | |
| Transfer multipole (V) | 4 | |
| Ion transfer target | High *m/z* | |
| Detector optimization | High *m/z* | |

**Table S1. Typical instrument parameters used for native MS and CD-MS measurements of rAAVs.**
